# Accurate and Fast feature selection workflow for high-dimensional omics data

**DOI:** 10.1101/144162

**Authors:** Yasset Perez-Riverol, Max Kun, Juan Antonio Vizcaíno, Marc-Phillip Hitz, Enrique Audain

**Affiliations:** European Molecular Biology Laboratory, European Bioinformatics Institute (EMBL-EBI), Wellcome Trust Genome Campus, Hinxton, Cambridge, CB10 1SD, UK.; RStudio, Boston MA; Department of Congenital Heart Disease and Pediatric Cardiology, Universitätsklinikum Schleswig–Holstein Kiel, Kiel, Germany.

**Keywords:** Feature Selection, R programming, Bioinformatics, Big Data, Proteomics, Genomics, Principal Component Analysis, Support Vector Machine

## Abstract

We are moving into the age of ‘Big Data’ in biomedical research and bioinformatics. This trend could be encapsulated in this simple formula: D = S × F, where the volume of data generated (D) increases in both dimensions: the number of samples (S) and the number of sample features (F). Frequently, a typical bioinformatics problem (e.g. classification) includes redundant and irrelevant features that can result, in the worst-case scenario, in false positive results. Then, Feature Selection (FS) constitutes an enormous challenge. Despite the number and diversity of algorithms available, the proper choice of an approach for facing a specific problem often falls in a ‘grey zone’. In this study, we select a subset of FS methods to develop an efficient workflow and an R package for bioinformatics machine learning problems. We cover relevant issues concerning FS, ranging from domain’s problems to algorithm solutions and computational tools. Finally, we use seven different proteomics and gene expression datasets to evaluate the workflow and guide the FS process.

## INTRODUCTION

The term ‘Big Data’ is often used to describe the huge volumes of information produced by modern systems such as mobile devices, tracking tools and sensors [1, 2]. In biomedical research, the growth of high-throughput (omics) technologies has resulted in an exponential growth in the dimensionality and sample size. This increase has two major directions: i) the number of samples processed, powered by novels machines (i.e. sequencers and mass spectrometers); and ii) the features, attributes and variables collected alongside each sample [3]. This high-dimensional environment becomes a challenge to many modelling tasks used in bioinformatics, ranging from sequence analysis to spectral analyses as well as literature mining. Reducing data complexity is therefore crucial for data analysis tasks, knowledge inference using machine learning algorithms, and data visualization [4–6].

The ‘curse of dimensionality’ (term first introduced by Bellman in 1957) [7] described the problem caused by the exponential increase in volume associated with adding extra dimensions to an Euclidean space. In this context, the typical bioinformatics problem involves both: relevant and redundant features. Therefore, a Feature Selection (FS) approach becomes a crucial and non-trivial task because: i) it provides a deeper insight into the underlying processes that are the foundation of the data; ii) it improves the performance (CPU-time and memory) of the machine learning (ML) step, by reducing the number of variables; and iii) it produces better model results avoiding overfitting. However, a FS algorithm brings an important decision in any machine learning workflow (e.g. classification of protein/gene expression profiles): are there redundant features (e.g. proteins or genes) in the dataset that are irrelevant and/or redundant for the biological study?

The most-common attempt to address the FS problem (the so-called univariate filtering approach) is to use a variable ranking method to filter out the least promising variables before using a multivariate method [8]. These methods have been used extensively in computational biology for cancer classification using microarray data [9, 10]. However, correlation filters could prompt some loss of relevant features that are meaningless by themselves but that can be useful in combination. To overcome this effect, a set of algorithms has been proposed to combine the original variables into a new and smaller subset of features, such as Principal Component Analysis (PCA) and Linear Discriminant Analysis. In PCA [11], new orthogonal features (latent variables or principal components) are obtained by maximizing the variation of the original features. The number of the latent features (factors) can be much lower than the number of original features, so that the data can be visualized in a much lower-dimensional space. As correlation filters, PCA methods can reduce the number of variables by looking into the feature dependencies without taking into account the final learning model. In 1997, a powerful strategy emerged that combines a FS algorithm with a learning/classification step: the so-called *wrapper* methods [12]. These wrapper approaches (e.g. *forward selection* and *backward elimination*) can use the prediction performance of a given machine learning approach to assess the relative usefulness of different subsets of variables. An exhaustive search can be performed if the number of variables is not too large.

Due to the diversity of FS methods available, it is hard to choose the correct approach needed to accomplish a specific task beforehand (e.g. regression or classification). In 2007, Saeys and co-workers published an introduction to FS in bioinformatics [3]. Also, several reviews have focused on the application in computational biology of particular methods such as PCA [13, 14] or Support Vector Machines (SVM) [15]. However, most of this work has been done to describe current methods in isolation and not to evaluate how they could be combined. In this manuscript, we developed a FS workflow and an R package for high-dimensional omics data analysis. The workflow combined univariate/multivariate correlation filters with wrapper feature backward elimination and it was applied to regression and classification problems. We benchmarked the individual steps of the described workflow, highlighting the optimal steps in different scenarios, using seven different omics datasets. Finally, we discuss major challenges when applying the described workflow to classification problems of high-dimensional omics data.

## MATERIALS AND METHODS

### Transcriptomic dataset of breast tumor samples (Dataset 1)

We first used a gene expression dataset (GEO (Gene Expression Omnibus) accession number: GSE5325) from *Saal et al*. [16], which has already been extensively studied before [13]. The authors performed a study using microarrays to measure the expression of 27,648 genes in 105 breast tumor samples. The dataset includes the estrogen receptor alpha status (0=negative, 1=positive), a transcription factor recognized as being important for stimulating the growth of a large proportion of breast cancers and used to explore co-expression [17].

### High-resolution isoelectric focusing proteomics dataset (Dataset 2)

The second dataset is the result of an electrophoresis experiment on peptide samples [18]. A total of 7,391 peptides were identified in 12 fractions, where each fraction corresponded to an experimental isoelectric point. This dataset has been used before to develop a machine learning model that can accurately predict the theoretical isoelectric points for peptides and proteins based on the amino acid sequence properties [5, 19].

### Triple-Negative Breast Cancer (TNBC) dataset (Dataset 3)

A third dataset containing protein quantification data using a label free technique was included [20]. The dataset assembles a panel of 44 (including samples and technical replicates) human breast cell lines and clinical tumors for analyzing the proteomics landscape of TNBC. The studied cell lines cover mesenchymal-, luminal-, and basal-like subtypes, as well as three receptor-positive and one non-tumorigenic cell lines. Thus, the idea behind including this dataset was to evaluate the ability of the proposed FS workflow to classify subtypes of cellular lines.

### Transcriptomics analysis of left ventricles of mouse hearts (Dataset 4)

A fourth dataset included the results of a transcriptomics analysis of left ventricles of mouse hearts subjected to an isoproterenol challenge [21]. In the study, the authors utilized expression arrays from left ventricular (LV) tissues, with and without an isoproterenol treatment, to understand the genetic control of gene expression and its relationship with heart failure. Then, the issue arising here suggests a binary classification problem where the researcher could be interested in, in order to know the optimal feature subset which could best discriminate between both classes (treated and non-treated samples).

### Expression data from normal and prostate tumor tissues (Datasets 5, 6, and 7)

Recently, Li *et al*. have used several gene expression datasets to benchmark different FS algorithms [22]. From the original microarray datasets, we have selected three of those datasets (GEO accession number: GSE6919), to compare the FS workflow with the results obtained by *Li et al*. **Note 1 (Supplemental Information 1)** summarizes the main characteristics of the datasets described previously.

## RESULTS AND DISCUSSION

A good feature subset can be defined as one that contains features highly correlated with (predictive of) outcome, yet uncorrelated (independent) with (not predictive of) each other. Nevertheless, the existing diversity of FS methods makes it challenging to choose the correct one for the task at hand (**Supplementary Information 1, Note 2**). Figure 1 represents the proposed overall workflow to perform FS in high-dimensional omics big data. First, a univariate correlation filter can be used before applying any wrapper approach, to determine the relation between each feature and the class or predicted variable. Then, a second filtering step (Correlation Matrix (CM) or PCA), can follow, in order to determine the dependencies between the different dataset features. Finally, backward elimination is achieved by wrapping a machine learning (ML) method, such as Random Forest and SVM around each example.

**Figure 1:**
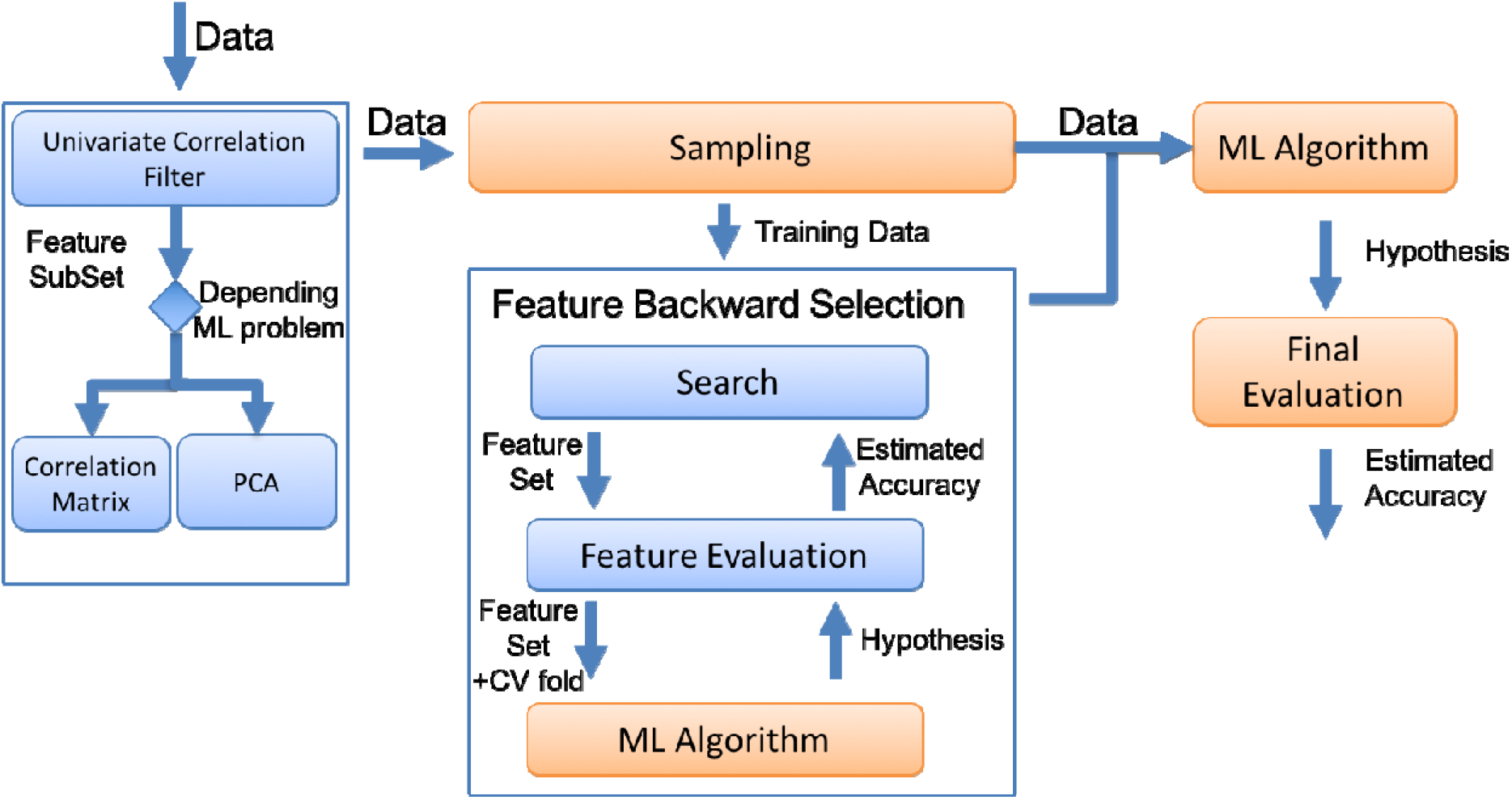
Proposed workflow for FS including a filtering step with univariate and/or multivariate approaches, followed by a wrapper approach (recursive feature elimination).

### Workflow R-package

An R-package has been developed to reproduce the proposed workflow (<u>https://github.com/enriquea/feseR</u>). For its development five main R packages were used: i) **Caret** [23] (**C**lassification **A**nd **RE**gression **T**raining) (<u>http://topepo.github.io/caret</u>), containing a set of functions that attempt to streamline the process for creating predictive models; ii) **randomForest** [24] is a package enabling Random Forest analysis (<u>https://cran.r-project.org/web/packages/randomForest/</u>); iii) **prcomp** is a native function included in the R package *stats*; iv) ***Kernlab*** [25] (<u>https://cran.r-project.org/web/packages/kernlab/</u>) provides the user with basic kernel functionality (e.g., computing a kernel matrix), along with some utility functions, commonly used in kernel-based methods; and v) the **FSelector** package [26] (<u>https://cran.r-project.org/web/packages/FSelector/</u>), which offers algorithms for filtering attributes (e.g. chi-squared, information gain, and linear correlation).

We have used the current FS workflow and R-package in combination two different machine learning (regression/classification) problems. Six of the datasets represent classification of (protein/gene) expression profiles and the last one a regression problem for the accurate estimation of isoelectric point of peptides and proteins. In the following sections, we discuss the results of combining the different steps of the FS workflow depending of the ML problem.

### Removing irrelevant features: Univariate correlation filtering

The univariate correlation filtering step removes all features that are not directly related to their class variables. When we applied this approach to **Dataset 1** it removed those genes with a non-correlated expression to the presence or absence of estrogen receptor alpha, reducing the number of genes from 8,534 (only those genes showing expression in all samples were considered) to 1,697. In **Dataset 2**, we used the univariate filter to remove features (amino acid properties) unrelated with the isoelectric point. Figure 2a shows the high-correlation found among the original 545 physicochemical peptides properties considered for the 7,391 peptides. We implemented a univariate correlation filter to remove all features that were not correlated with the isoelectric point, reducing the number of variables to 81 features (Figure 2b). When we extended the analysis to the remaining benchmarking datasets, we observed that, in general, univariate correlation filtering removed more than 80% of the original features that were not related to the predicted variable. As previously discussed by other authors [8], univariate correlation filtering should be always applied at early stages of any classification and/or regression process. However, univariate correlation filtering can only be used to study the relationship of each feature with a class variable, but cannot be applied to find the relationships among them. For this reason, a multivariate step (e.g. correlation matrix) was used (Figure 1) to remove the redundancy among highly-correlated features.

**Figure 2:**
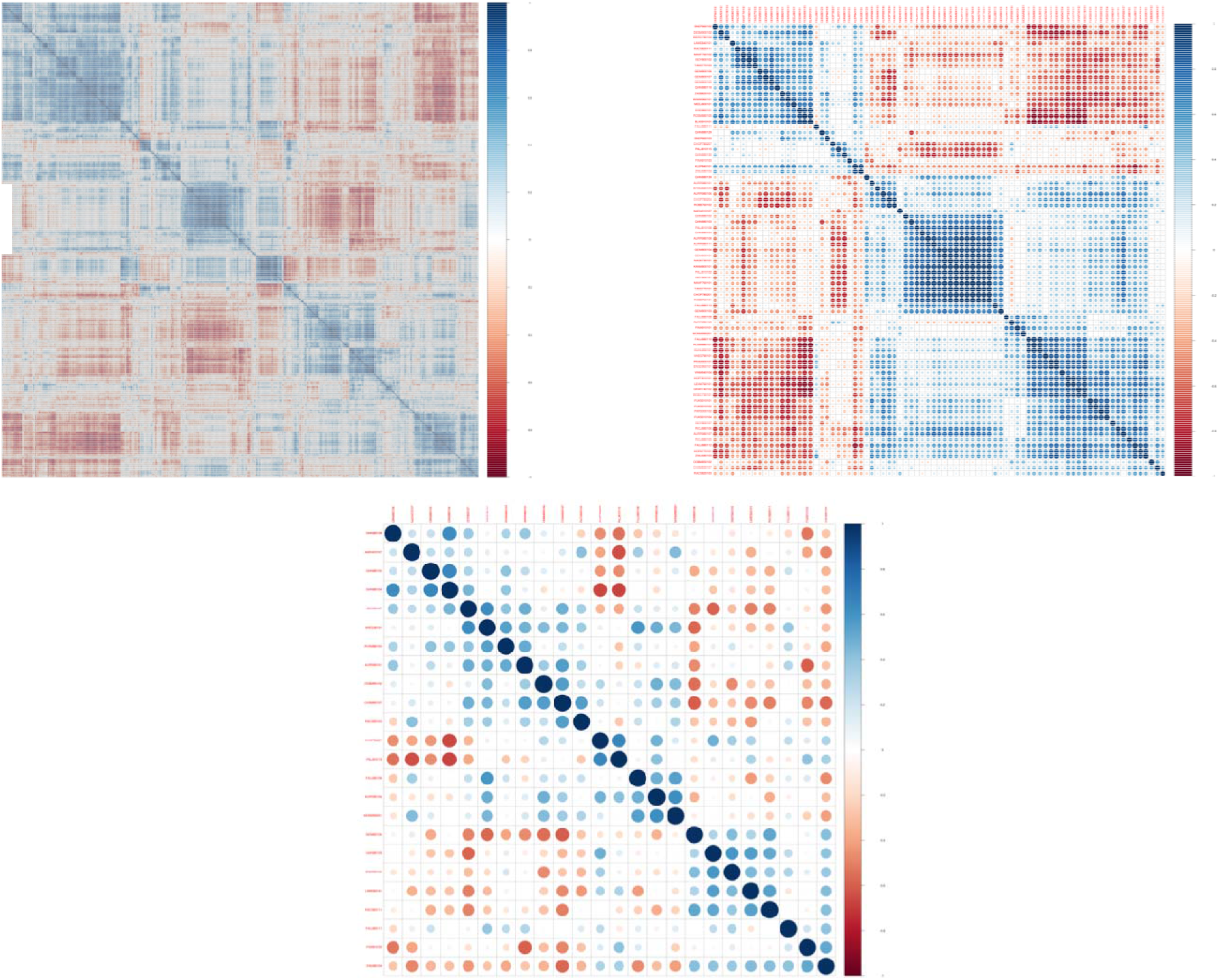
(**a**) Correlation matrix for the 544 physicochemical (features) of the 7,391 peptides (samples) included in Dataset 2; (**b**) 81 relevant features after univariate correlation filtering; (**c**) the final 21 variables after the correlation-matrix filtering steps.

### Reducing feature Complexity: CM or PCA

We implemented two different strategies (depending on the classification or regression problem) to reduce the number of variables, while keeping most of the original and relevant information: CM and PCA. **Dataset 2** is a good example of a dataset containing regression related problems. In this particular case, the aim was to predict more accurately the isoelectric point of peptides and proteins, using other physicochemical features of the peptides. Therefore, the final model should be based on, or be correlated to, the original features (because they would be used in the future to make a predictor that could be applied for other datasets). One of the simplest and most powerful filtering approaches to remove feature redundancy, while keeping original features, is the use of a CM filter. For example, peptides properties such as aromatic rings, bond and carbon atom counts are strongly correlated [5, 27]. Therefore, any of these variables could be used as a proxy for all the others. Figure 2b shows the correlation matrix of the feature space after the univariate filtering step (81 variables). It should be noted that several features clustered together, suggesting a high-redundancy in the feature set. By applying the CM filter, it is possible to remove those that are redundant (or irrelevant) and to keep only a reduced feature set for subsequent analysis steps (Figure 2c). The present workflow keeps only 21 variables (out of the original 545 features) for the final machine learning step (Figure 1). The current approach also reuses the final model in new datasets because the filtering steps preserve the original variables by only removing the redundant ones.

Opposite to **Dataset 2**, the other datasets constitute good examples of classification related problems. In addition to the MC filter approach, we implemented and studied the use of Principal Component Analysis (PCA) as a multivariate filter to reduce the number of features. PCA reduces the dimensionality of the data while retaining most of the variation in the predictor variables [13]. Thus, by using a few components, PCA can represent each sample by using relatively few (new) variables instead of (potentially) thousands of them. Figure 3 shows the PCA performed in **Dataset 1**. The proportion of the variation present in all genes is encompassed within each of the principal components, with the first few components representing most of it (Figure 3a). The cumulative variance analysis shows that most of the variance is contained in the first 30 principal components (75%), where only 76 components reach a 95% of variance (Figure 3b) and 104 components are enough to retain all the original variance. This number of variables is 10-fold smaller than the original 1,697 features obtained after applying the univariate correlation filter.

**Figure 3.**
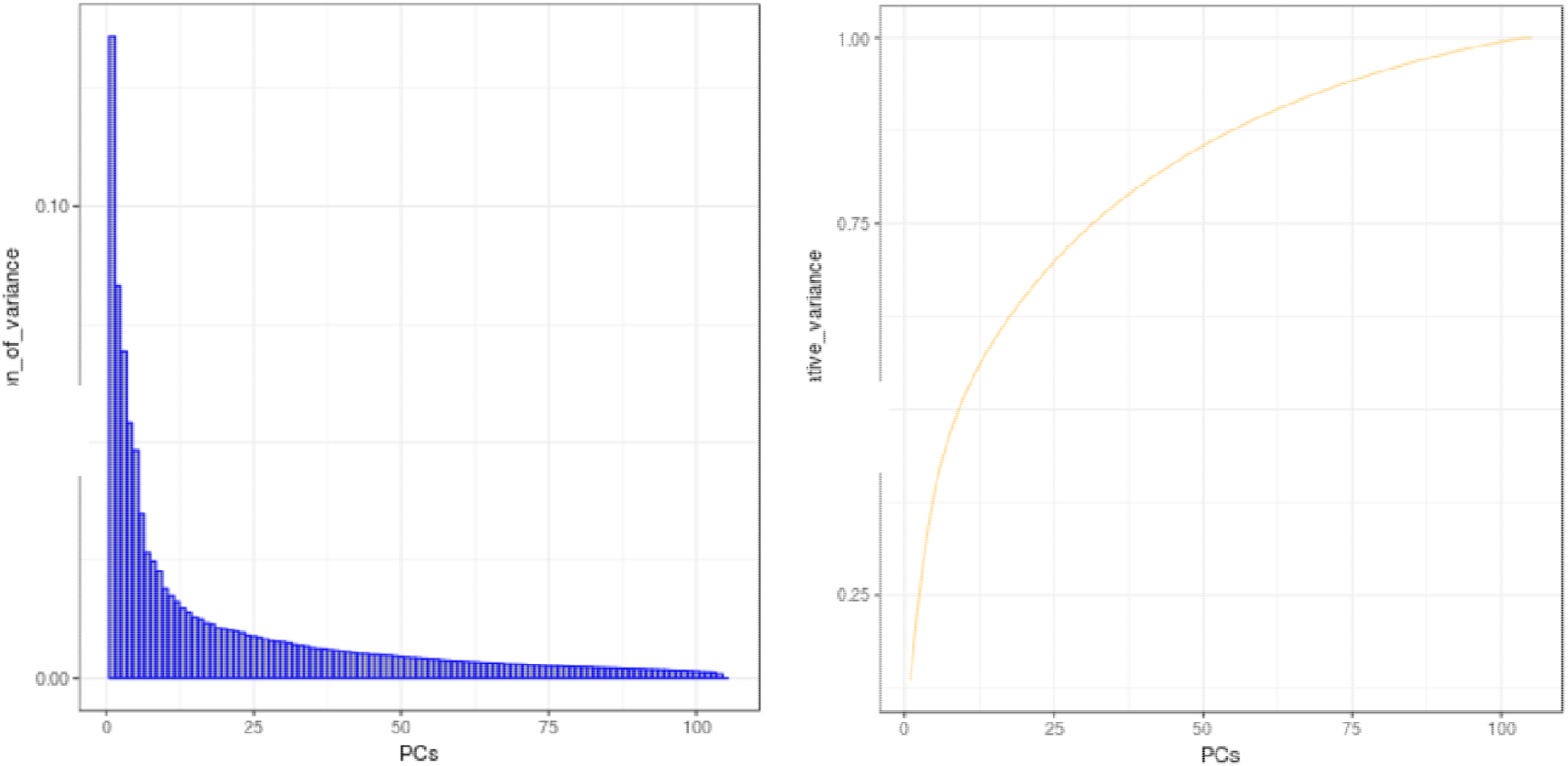
(**a**) Proportion of variance and (**b**) cumulative variance of principal components for the analysis of **Dataset 1**.

When the number of variables is larger than the number of samples, PCA can reduce the dimensionality of the samples to, at most, the number of samples, without losing information [13, 28]. We obtained the same results when PCA was applied to the other relevant datasets (**Dataset 3** to **Dataset 7**, those with a classification problem, **Supplementary Information 2**). However, since the principal components are linear combinations of the original data, it is not obvious how model parameter estimates can relate back to the original variables. Thus, this method is not suitable for problems where it is required to keep the primary information (e.g. in the case of regression problems, **Dataset 2**).

### Optimizing the Feature selection: Wrapper Recursive Feature Elimination

All filtering FS approaches previously shown (e.g. correlation-based or PCA) are relatively easy to implement and computationally fast. Therefore, these algorithms represent a suitable choice in the first stage of any given FS pipeline. However, wrapper methods should be used in the last steps to find the “optimal” feature subsets, by iteratively selecting features based on classifier performance (Figure 1). The wrapper methods should be combined with cross-validation steps to improve the final results [12, 29]. These cross-validation steps can be used to assess the results of the learning analysis (e.g. regression or classification) and help to generalize these steps to an independent dataset. The goal of cross-validation is to define a dataset to “test” the model in the training phase (i.e., the validation dataset), in order to limit problems like overfitting [29]. In the proposed workflow, we used a recursive feature elimination (backward elimination) approach in combination with two machine learning models (Random Forest and SVM) to systematically increase each machine learning step. The number of cross-validation iterations should be evaluated in detail because it could significantly increase the running time without improving the performance of the model prediction.

We implemented the wrapper backward elimination step in combination with the SVM radial kernel, in order to predict the isoelectric point using **Dataset 2.** Table 1 shows the performance (regarding running time and model prediction accuracy) of the feature workflow for **Dataset 2**. We benchmarked all the FS combinations with the SVM model by removing each of them. Applying the SVM model alone (**SVM**) without FS or cross-validation helps to predict the isoelectric point with a high root-mean-square error (RMSE) of 0.88. In contrast, when both correlation filters (**X2-CM-SVM**) were applied, RMSE and running time decreased to 0.57 and 0.50 min, respectively. When the complete workflow (**X2-CM-RFE-SVM-CV3**) was used RMSE decreased to 0.33 min (Table 1). It should be noted that when pre-filtering was applied (**RFE-SVM-CV3**), RMSE decreased to 0.32 and two new variables were added to the SVM model. However, this improvement in performance (e.g. low RMSE) decreased the overall efficiency of the workflow by increasing the execution time three-fold. Also, we observed no changes where the number of cross-validation steps was increased (Table 1).

**Table 1:**
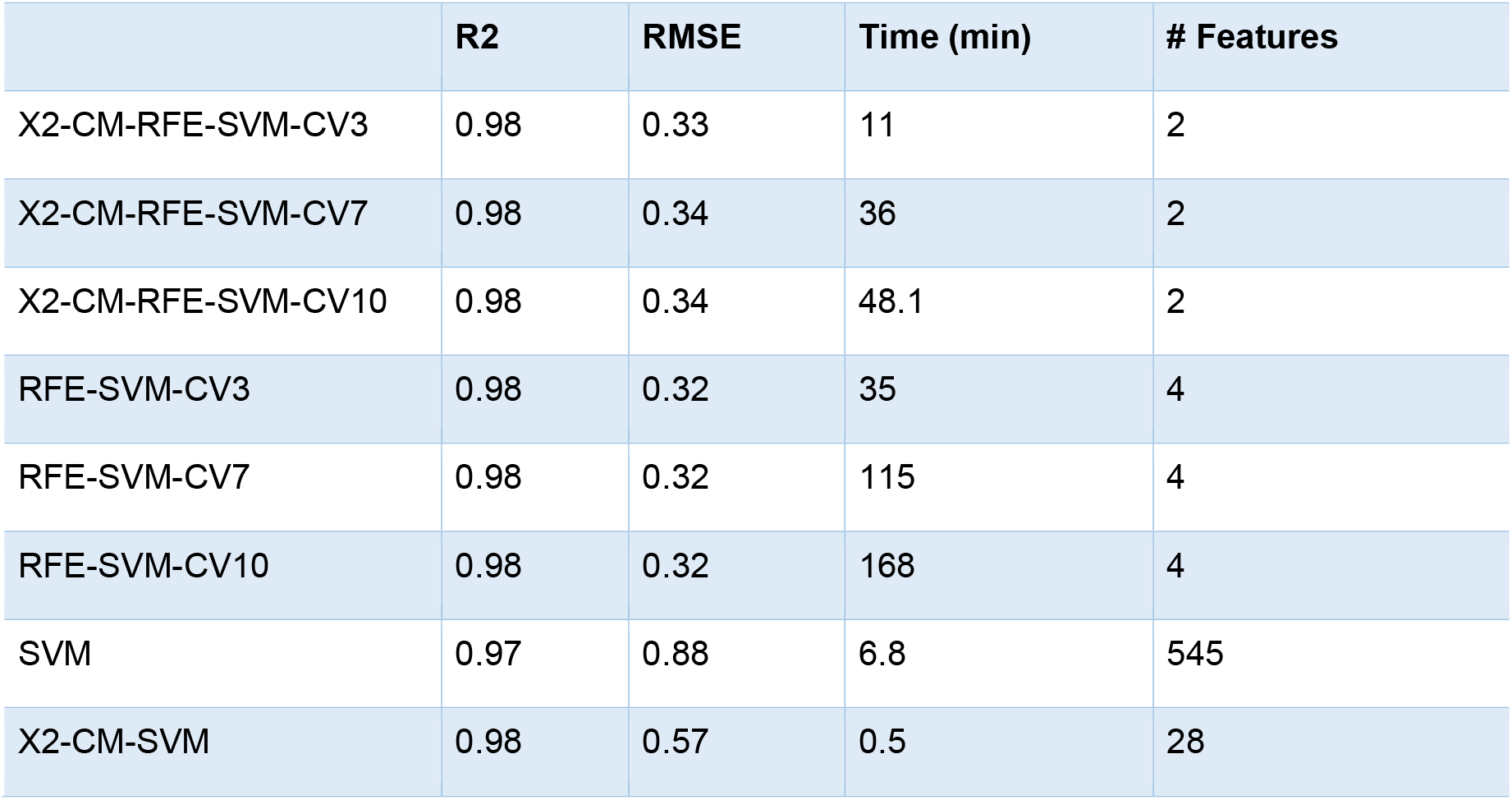
Benchmark of the SVM regression model for **Dataset 2** applying different FS methods (**SVM**), no feature selection, (**X2**) univariate correlation alone, (**CM**) correlation matrix filtering, (**RFE**) and wrapper feature elimination. The figures indicated using the prefixes CV3, CV7 and CV10 correspond to the number of interactions in the cross-validation steps during the RFE feature selection.

|  | **R2** | **RMSE** | **Time (min)** | **# Features** |
| --- | --- | --- | --- | --- |
| X2-CM-RFE-SVM-CV3 | 0.98 | 0.33 | 11 | 2 |
| X2-CM-RFE-SVM-CV7 | 0.98 | 0.34 | 36 | 2 |
| X2-CM-RFE-SVM-CV10 | 0.98 | 0.34 | 48.1 | 2 |
| RFE-SVM-CV3 | 0.98 | 0.32 | 35 | 4 |
| RFE-SVM-CV7 | 0.98 | 0.32 | 115 | 4 |
| RFE-SVM-CV10 | 0.98 | 0.32 | 168 | 4 |
| SVM | 0.97 | 0.88 | 6.8 | 545 |
| X2-CM-SVM | 0.98 | 0.57 | 0.5 | 28 |

Wrapper backward elimination step provided a powerful method to optimize the final subset of variables in response to the regression SVM model. Figure 4 shows the final results of the isoelectric point prediction (**Dataset 2**) for all FS combinations. Backward selection in combination with the cross-validation step enables a better estimation of the variable prediction (isoelectric point) in the regions where less experimental evidences exist (basic pH range). This workflow has been used in a recent approach to predict the isoelectric point and it has proven to predict the isoelectric point more accurately than any other algorithm so far. A similar implementation was applied to the remaining datasets (1,3-7) where a Random Forest model was wrapped around, using a recursive approach to evaluate the performance and the variable weight following different FS workflows. We first evaluated the Random Forest approach for FS without any filtering and parameter tuning as discussed before by *Díaz-Uriarte et al*. [30]. In addition, four recursive feature elimination methods, wrapped with Random Forest, were combined as follows: **RFE-RF** without any pre-filtering step (i.e. other FS methods), PCA combined with **RFE-RF**, univariate correlation filtering (**X2**) combined with **RFE-RF**, and finally, all methods were used sequentially: **X2-PCA-RFE-RF** or **X2-MC-RFE-RF**

**Figure 4:**
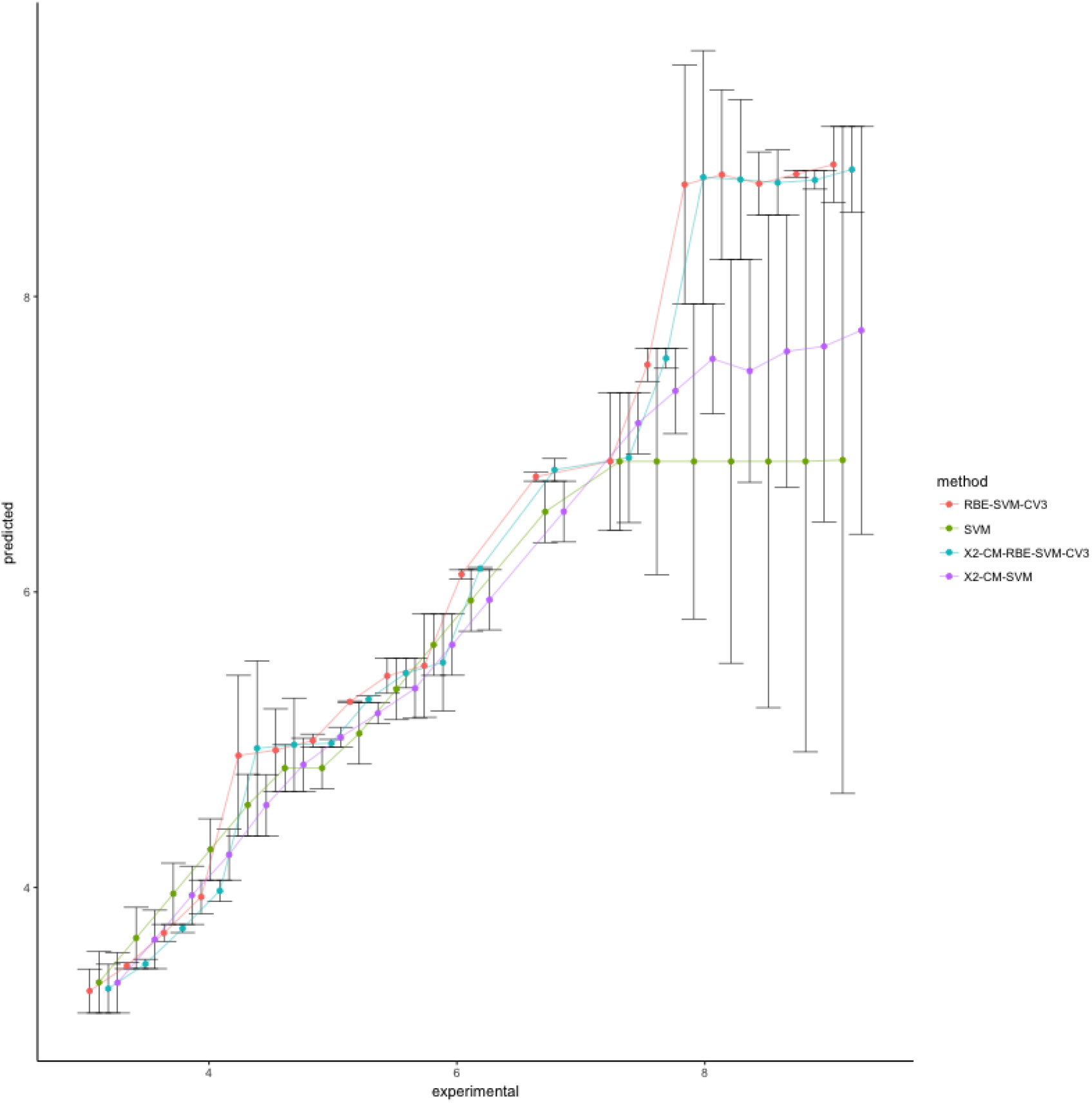
Error plot of predicted isoelectric point vs the experimental isoelectric point (**Dataset 2**): (**SVM**) applying FS or cross-correlation step; (**X2-CM-SVM**) adding correlation filters as the only steps for feature selection; (**RFE-SVM-CV3**) recursive feature elimination, three interactions of cross-validation combined with **SVM**; (**X2-CM-RFE-SVM-CV3**) considering the full FS workflow.

Figure 5 shows the performance evaluation (for the expression datasets 1, 3-7) of each complete FS combination (**X2-PCA-RFE-RF** and **X2-MC-RFE-RF**) and the random forest classification without FS step. We use the approach previously reported by *Pochet et al*. [31], where 20-fold randomized test data were used to summarize the accuracy in the prediction (see detailed description in **Supplementary Information 2**). Also, we kept a 10-fold internal crossvalidation step in all implementations of recursive feature selection trials. The results shown that when any of the full FS approaches are applied the average accuracy is higher compare with the results when not FS is used (red box plots). Only, in **Dataset 3** the workflow using PCA is less efficient than the random forest without FS step which can be related with the low number of samples analyzed (44). Importantly, even when RF perform very well it retains all the original features on each making difficult to decided which features are more relevant for the classification (**Supplementary Information 1**, Table 2). Both FS workflows reduce the number of variables in all cases in more than 90% (**Supplementary Information 1**, Table 2), with average accuracy always above 70% (Figure 5). Because both workflow shows similar performance and some users may want to select PCA (less variables) or MC (original features), the R-package allows to define which multivariate option use during the FS.

**Figure 5:**
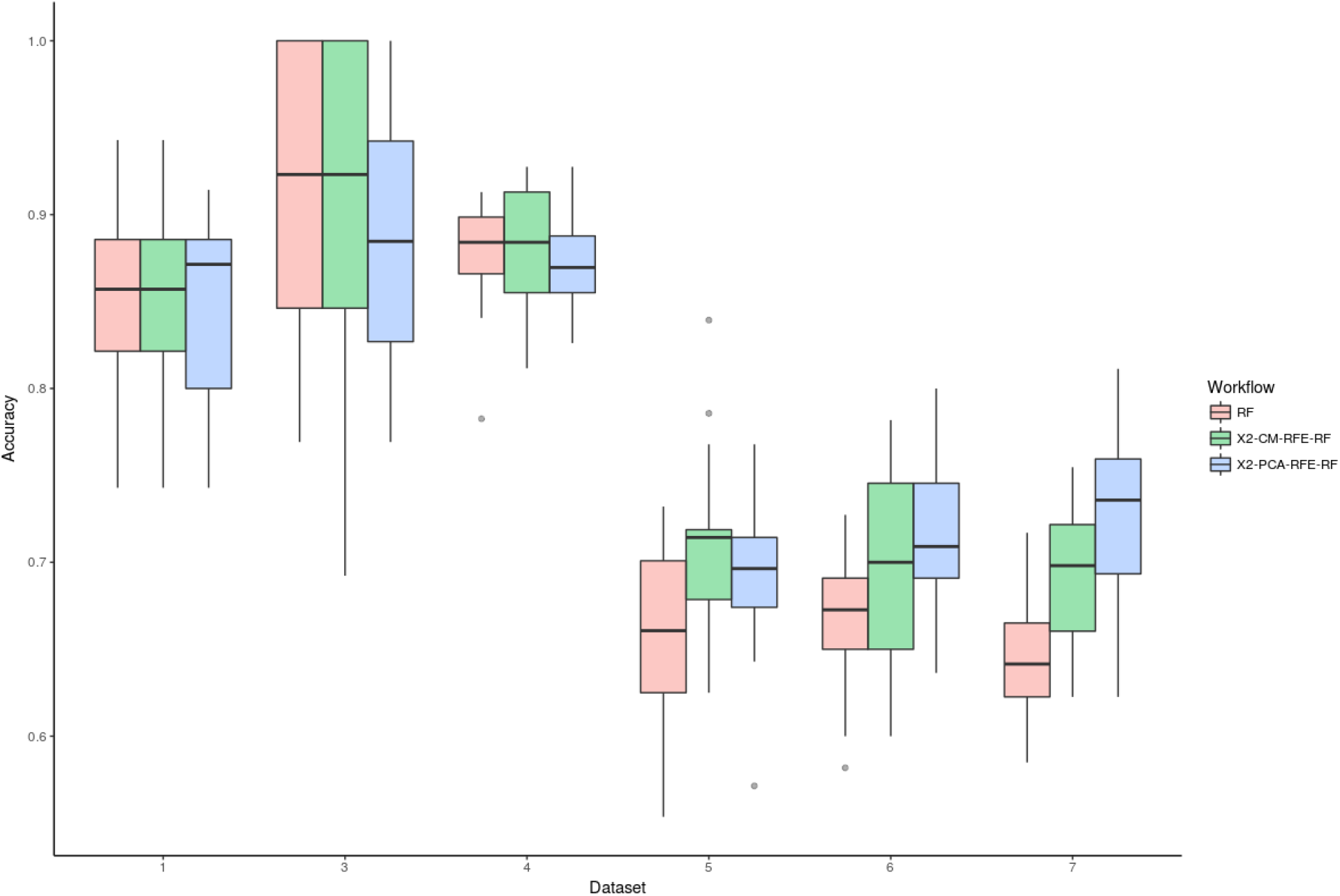
Accuracy vs. Feature Selection combination for Expression datasets (1, 3, 4, 5, 6 and 7). (**RF**) Random Forest without previous feature selection step; (**X2-CM-RFE-RF**), random forest classification after the feature selection step using univariate correlation filter with matrix correlation and recursive feature elimination; (**X2-PCA-RFE-RF**), random forest classification after the feature selection step using univariate correlation filter with principal component analysis and recursive feature elimination. All methods include an internal cross-validation 10-fold step. All accuracy metrics were estimated following the approach previously reported by *Pochet et al*. [31], where 20-fold randomized test data were used to summarize the accuracy of the FS combination.

**Table 2:** Benchmarking of the Random Forest model (classification) for **Dataset 1**, when different FS methods are applied: (**RF**) Random Forest only, (**RFE**) wrapper recursive feature elimination with 10-times internal cross-validation, (**PCA**) Principal Component Analysis with (**X2**) univariate correlation filtering. Each method is applied 20 times with randomized and class-balanced training datasets. The accuracy values provided correspond to the average value.

|  | Accuracy (%) | SD | Time (min) | # features |
| --- | --- | --- | --- | --- |
| RF | 83.46 | 8.1 | 1.46 | 1969 |
| RFE-RF | 84.61 | 6.3 | 15.83 | 30 |
| PCA-RFE-RF | 83.43 | 5.4 | 3.12 | 10 |
| X2-RFE-RF | 87.04 | 3.6 | 4.92 | 25 |
| X2-PCA-RFE-RF | 88.21 | 4.5 | 3.51 | 8 |
| X2-MC-RFE-RF | 85.01 | 5.7 | 6.35 | 8 |

Table 2 summarizes the benchmark metrics (accuracy, standard deviation, number of final features and time) for each evaluated FS workflow (in **Dataset 1**). While all methods kept the accuracy in the range 83-88%, when all methods were combined (proposed workflow) a lower standard deviation was obtained. Using a Random Forest model without FS, the classification process was faster than in the case of any other combination, keeping all the relevant features (1,969 of them). Including PCA and Recursive Feature Elimination (**PCA-RFE-RF**), we observed a strong feature reduction (7-10 components) and a better standard deviation (5.4). Selecting a univariate correlation filter (**X2-RFE-RF**), a lowest standard deviation was obtained (3.2).

Figure 6 visualizes the results of the Random Forest classification algorithm without (panels a, c, f) and with (panels b, d, e) a FS step; for Datasets 1, 3, and 4, respectively. The results show that the remaining features obtained allow to ‘discriminate’ between the different samples classes or groups (see detailed description in **Supplementary Information 2**). It can be concluded that for those classification problems where the original features are needed, the PCA step could be removed without sacrificing general performance (accuracy, standard deviation, or CPU time). In contrast, univariate correlation filtering FS steps had a key impact on the final results of the Random Forest model by increasing the performance in all the studied combinations. As we pointed out earlier, PCA ‘obfuscates’ the primary information, and thus, can potentially result in problems. When it is desirable to keep the “initial nature” of the variables, filtering methods (e.g. univariate correlation filter) exhibit a good performance (**Table 1-2**) with a considerable lower number of features (25 final features).

**Figure 6:**
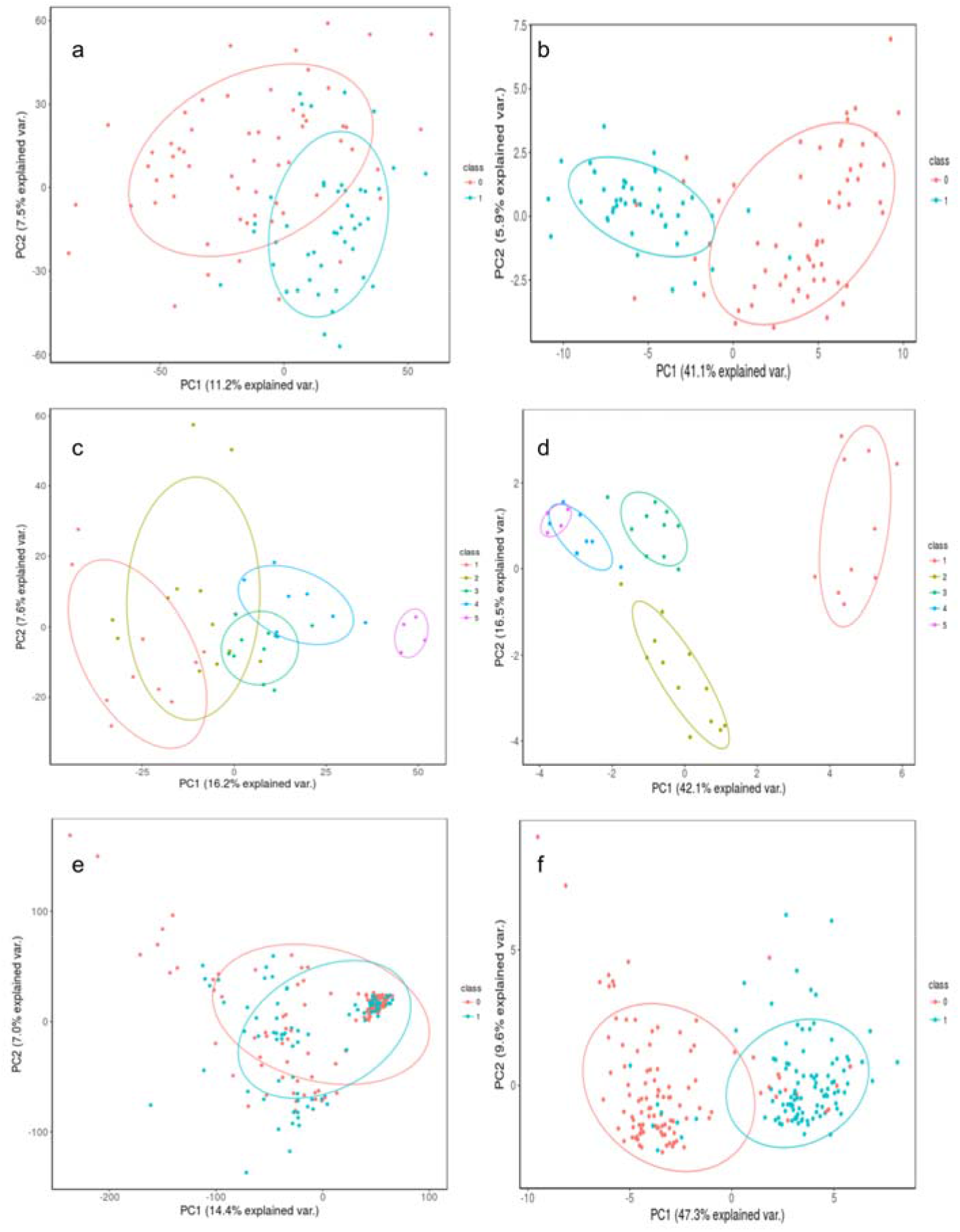
Visualization of the classification process using the first two Principal Components (PC1 and PC2) from the original data before (panels a, c, and e) and after (panels b, d, and f), to apply the following FS workflow: Univariate correlation (X2) with Recursive Feature Elimination (RFE) wrapped with Random Forest (RF). The figure shows the classes distribution for **Dataset 1** (panels a-b), **Dataset 3** (panels c-d) and **Dataset 4** (panels e-f).

### Summary of the benchmarking process

We have demonstrated the impact of the FS workflow in the classification and/or regression results as well as in the performance of the machine learning algorithm (CPU time and memory). Finally, we applied the same FS workflow to gene expression data from normal and prostate tumor tissues (**Datasets 5, 6** and **7**), and compared them with the results obtained by *Li et al*. [22], who used a similar approach on the same datasets (see Table 9 in [22]). Even though we observe a slight improvement in the classification accuracy in these three datasets (Table 3), the most notable differences were found in the number of features obtained by the final models and in the total runtime, using a similar computational platform. Thus, the results from the comparison reinforce our previous observations and validate the effectiveness of the FS workflow proposed in this manuscript.

**Table 3:** Performance comparison between the proposed approach (**X2-PCA-RFE-RF**) and the method reported by Li *et al*. [22]. The computer used in the original manuscript was an Intel(R) Core(TM) i5-4690 @ 3.5 GHz CPU, with 16 GB of RAM. In this study, we used an Intel(R) Core(TM) i5-4200 @ 2.5 GHz CPU, with 16 GB of RAM.

| Dataset | Method | Accuracy | Variables | Runtime (min) |
| --- | --- | --- | --- | --- |
| **GSE6919/GPL8300** | Current Workflow | 0.77 | 35 | 8.50 |
|  | Li *et al.* | 0.72 | 92 | 74.30 |
| **GSE6919/GPL92** | Current Workflow | 0.80 | 5 | 9.11 |
|  | Li *et al.* | 0.73 | 174 | 71.50 |
| **GSE6919/GPL93** | Current Workflow | 0.81 | 6 | 12.00 |
|  | Li *et al.* | 0.71 | 121 | 68.60 |

### Conclusions

FS selection algorithms are playing a major role to select correct variables for different classification and regression problems. Nevertheless, choosing the appropriate algorithm (or combination of algorithms) is not a trivial task. Different studies have highlighted methods to perform FS, but unfortunately, a thorough comparison including proper benchmarking is still lacking. Another major challenge remains: how to efficiently combine different FS methods to improve the final results. The developed FS workflow shown in this manuscript combines major strengths of univariate filtering methods, with CM and PCA strategies, as well as recursive feature elimination in two well-known learning problems: classification and regression. When univariate filtering was used in both types of problems the number of features was reduced by 80% without compromising the accuracy of the final model, and decreasing the CPU time of the learning model steps. The introduction of a wrapper method (recursive feature elimination) in combination with the learning model improved the accuracy in both cases. If the wrapper method is applied without a previous filtering step, the CPU-time becomes too high. Finally, we demonstrated that the use of an intermediate FS step to remove redundancy between variables and features can significantly increase the accuracy of the learning model. This can be achieved by transforming the original variables into new components (retaining most of the variability in the original values) using PCA or by removing redundant highly correlated variables.

Large efforts have taken place in recent years to adopt individual FS methods. However, in our opinion, a multiple FS step workflow offers more promising results. Future developments should focus on other fields where the number of samples is growing considerably (e.g. clinical genomics, text and literature mining), and on the combination of heterogeneous datasets from different sources.

## Acknowledgements

Y.P.-R. is supported by BBSRC ‘PROCESS’ grant (BB/K01997X/1). J.A.V. acknowledges the Wellcome Trust (grant number WT101477MA) and EMBL core funding.

## Abbreviations

CM: Correlation Matrix
FS: Feature Selection
ML: Machine Learning
PCA: Principal Component Analysis
RFE: Recursive Feature Elimination
RMSE: Root Mean Square Error
RF: Random Forest
SVM: Support Vector Machine
TNBC: Triple-Negative Breast Cancer
X2: Univariate Correlation

